# Demonstration of End-to-End Automation of DNA Data Storage

**DOI:** 10.1101/439521

**Authors:** Christopher N. Takahashi, Bichlien H. Nguyen, Karin Strauss, Luis Ceze

**Affiliations:** School of Computer Science and Engineering, University of Washington; Microsoft Research

## Abstract

We developed a complete end-to-end DNA data storage device. The device enables the encoding of data, which is then written to a DNA oligonucleotide using a custom DNA synthesizer, pooled for liquid storage, and read using a nanopore sequencer and a novel, minimal preparation protocol. We demonstrate an automated 5-byte write, store, and read cycle with the ability to expand as new technology is available.

## Main

Storing information in DNA is an emerging technology with considerable potential to be the next generation storage medium of choice. Although contemporary approaches are book-ended with mostly automated synthesis[1] and sequencing technologies (e.g., column synthesis, chip array, ink-jet, Illumina, nanopore, etc.), significant intermediate steps remain largely manual [2, 3, 4]. Without full automation, DNA data storage is unlikely to become a more prevalent option for applications other than extreme archiving.

To demonstrate the practicality of integrating fluidics, electronics and infrastructure, and explore the challenges of full automation, we developed the first full end-to-end automated DNA storage device. Our device is intended to act as a proof of concept that provides a foundation for continuous improvements as well as a first application for modules that can be used future molecular computing research. As such, we adhered to specific design principles for the implementation: (1) maximize modularity for the sake of replication and reuse, and (2) reduce system complexity to balance cost and the labor input required to setup and run the device modules.

Our resulting system has three core components that accomplish the write and read operations (Fig. 1a): an encode/decode software module, a DNA synthesis module, and a DNA preparation and sequencing module (Fig. 1b, c). It has a bench-top footprint and costs approximately $10k USD, though careful calibration and elimination of costly sensors and actuators could reduce the cost to approximately $3k-4k USD.

Before a file can be written to DNA its data must first be translated from 1’s and 0’s to A’s, C’s, T’s, and G’s. The **encode software module** is responsible for the translation (see methods and [5]) and the addition of error correction into the payload sequence. Once the payload sequence is generated, additional bases are added to ensure its primary and secondary structure is compatible with the read process and the DNA sequence is sent to the synthesis module for instantiation into physical DNA.

The **synthesis module** is built around two valved manifolds that separately deliver hydrous and anhydrous reagents to the synthesis column. Our initial designs used standard valves, but the dead volume at junction points caused unacceptable contamination between cycles. Therefore, we switched to zero dead volume valves [6]. The combined flow path is then monitored by a flow sensor, whose output is coupled to a standard fitting; the fitting can be coupled to arbitrary devices, such as a flow cell for chip array synthesis [7] or, in this case, adapted to fit a standard synthesis column. Once synthesis is complete, the synthesized DNA is eluted into a storage vessel, where it is stored until retrieval.

When a read operation is requested, the stored DNA pool’s volume is reduced to 2 μL to 4 μL by discarding excess DNA through the waste port. A syringe pump in the **DNA preparation and sequencing module** then dispenses our single-step preparation/sequencing mix (Fig. 1d) into the storage vessel; positive pressure pushes the mixture into the ONT MinION’s priming port (Fig. 1[b, c]-3). We chose the MinION as our sequencing device due to its low cost, ease of automation, and high throughput. However, it is not capable of reading unmodified DNA, nor is it optimized for reading short DNA oligonucleotides [8]. To mitigate these limitations, we developed a single-step MinION preparation protocol that requires only payload DNA and a master mix containing a customized adapter (Fig. 1d), T4 ligase, ATP, and a buffer. Each payload sequence is constructed to form a hairpin structure with a specific 5’ 4-base overhang. The customized adapter has a complementary overhang, which aids T4-mediated, sticky-ended ligation. To sequence, the payload and master mix are combined and incubated at room temperature for 30 minutes. Thereafter, the mixture is directly loaded into the MinION through the priming port. Since the introduction of air bubbles causes sequencing failure we built a 3D printed bubble detector that valves off the loading port immediately after detecting the gas that is aspirated following the sample. This allows us load nearly the full sample without damaging the flow cell. Additionally, while not demonstrated here, other research suggests that random access via selective ligation over a small set of sequence identifiers (≈ 20) can be achieved using orthogonal sticky ends during preparation [9].

**Figure 1:**
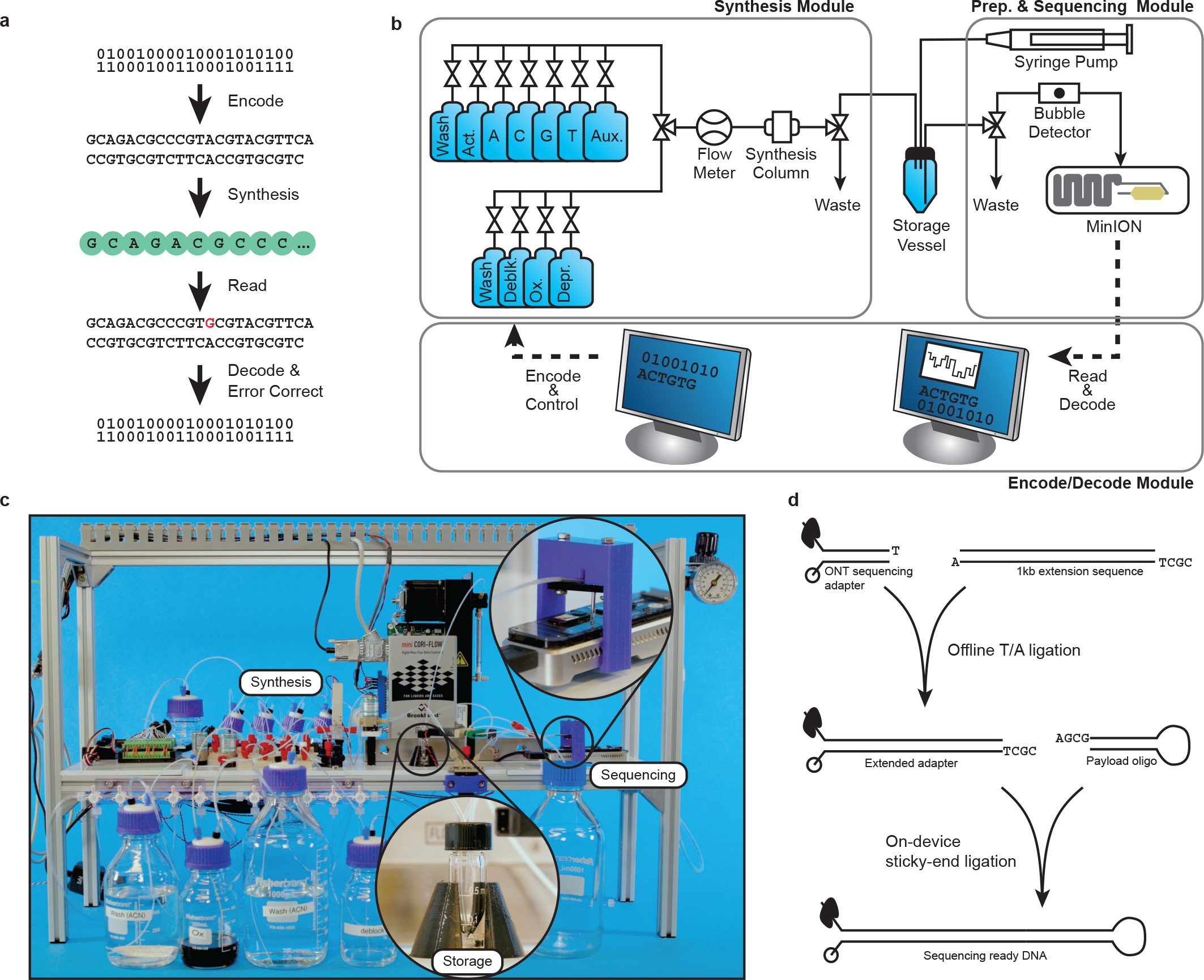
An overview of the write-store-read process. Data is encoded, with error correction, into DNA bases, which are synthesized into physical DNA molecules and stored. When a user wishes to read the data, the stored DNA is read by a DNA sequencer into bases and software corrects any errors retrieving the original data. a) The logical flow from bits to bases to DNA and back. b) A block diagram representation of the system hardware’s three modules: synthesis, storage, and sequencing. c) A photograph showing the completed system. Highlighted are the storage vessel and the nanopore loading fixture. The majority of the remaining hardware is responsible for synthesis. d) Overview of enzymatic preparation for DNA sequencing. An arbitrary 1 kilobase “extension segment” of DNA is PCR amplified with TAQ polymerase, and a Bsa-I restriction site is added by the primer, leaving an A-tail and a TCGC sticky end after digestion. The extension segment is then T/A ligated to the standard Oxford Nanopore Technology (ONT) LSK-108 kit sequencing adapter, creating the “extended ONT adapter,” which ensures that sufficient bases are read for successful base calling. For sequencing, the payload hairpin and extended adapter are ligated, forming a sequence-ready construct that does not require purification.

Once sequencing begins, the **decode software module** aligns each read to the 1k base extension region and the poly-T hairpin. If the intervening region of DNA is the correct length, the decoder attempts to error check/correct the payload using a Hamming code with an additional parity bit; the code corrects all single-base errors and detects all double-base errors. Once the payload is successfully decoded, it is considered correct if it matches a 6-base hash contained with the data. At this point, sequencing terminates, and the MinION flow cell may be washed and stored for later reuse.

Our system’s write-to-read latency is approximately 21 h. The majority of this time is taken by synthesis, viz., approximately 305 s per base, or 8.4 h to synthesize a 99-mer payload and 12 h to cleave and deprotect the oligonucleotides at room temperature. After synthesis, preparation takes an additional 30 min, and nanopore reading and online decoding take 6 min.

**Figure 2:**
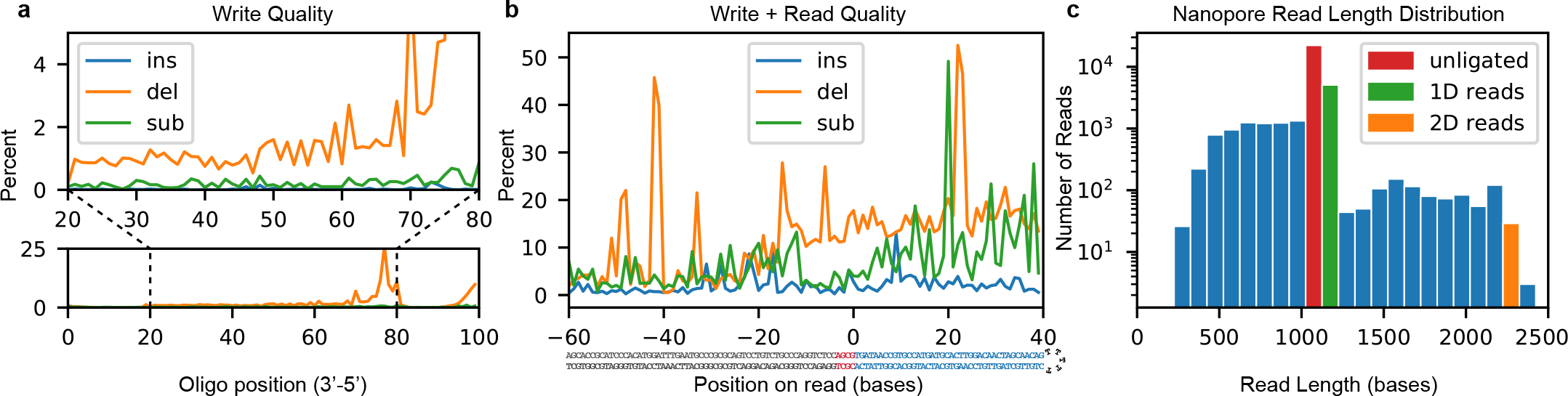
Synthesis and sequencing process quality. a) Insertion, deletion, and substitution frequency by locus for a synthesized and PCR-amplified 100-mer. Below: An overview of errors. Above: An expanded view of the central 60 bases. The terminal 20 bases come from primers used in amplification and therefore are not representative of synthesis quality. b) Combined write-to-read quality of synthesis, ligation, and sequencing. Bases −60 to −4 (below, grey) are adapter bases. Bases −3 to 0 (below, red) are the ligation scar. Bases 0 to 39 (below, blue) are the synthesized payload region with 8 bases of padding on the 3’ end. c) Distribution of nanopore read lengths with unligated, 1D and 2D read lengths identified.

Using this prototype system, we stored and subsequently retrieved the 5-byte message “`HELLO`” (`01001000 01000101 01001100 01001100 01001111` in bits). Synthesis yielded approximately 1 mg of DNA, with approximately 4 μg≈100 pmol retained for sequencing. Nanopore sequencing yielded 3469 reads, 1973 of which aligned to our adapter sequence. Of the aligned sequences, 30 had extractable payload regions. Of those, 1 was successfully decoded.

Inspecting the sequencing data indicates that the low payload yield was largely due to two factors. The first and primary factor is *low ligation efficiency*. Although chemical conditions should be optimal for T4 ligase, incomplete strands from the unpurified synthesis product likely outcompeted full-length strands, leading to a poor apparent ligation rate of less than 10% (Figure 2c). The second factor is *read and write fidelity*. To interrogate the write error rate, we synthesized a randomly generated 100-base oligonucleotide with distinct 5’ and 3’ primer sequences. The oligonucleotide was then PCR amplified and next-gen sequenced to reveal: an error rate of almost zero insertions; <1% substitutions; and 1-2% deletions (Fig. 2a) for most positions, with increased deletions toward the 5’ end due to increased steric hindrance as strand length increases [10]. Literature suggests a nanopore error rate near 10% [8, 11], so we also performed a synthesis-to-sequencing error rate analysis on an 89-mer hairpin sequence, encoding “`HELLO`” in its first 32 payload bases. Figure 2b shows the read error when aligned to the extended adapter and payload sequence. Bases −60 to −1 were directly PCR amplified from the lambda genome and given a good baseline for nanopore sequencing fidelity under our conditions; bases 0 through +40 come from the payload region and characterize the total write-to-read error rate. The complex combination of these errors — especially deletions and read truncations — causes many strands to be discarded before a decoding attempt is made.

We demonstrated the first fully automated end-to-end DNA data storage device. This device establishes a baseline from which new improvements may be made toward a device that eventually operates at a commercially viable scale and throughput. While 5 bytes in 21 hours is not yet commercially viable, there is precedent for many orders of magnitude improvement in data storage [12], and the underlying physics and chemistry show impressive upper bounds for density [4]. Furthermore, the modules and methods developed here are now being applied to other research projects internally with some publications forthcoming. Near-term improvements to this system will include more efficient sequencing preparation, optimized synthesis, and halving of write-to-read times through improved deprotection conditions [13]. Additionally, a cost-optimized version could be designed by eliminating the syringe pump and flow sensor, both unnecessary if flow rates are well measured and calibrated. This could save approximately 60% of our current device’s cost at the expense of a device that is more laborious to operate. Future improvements will focus on storage density, coding, and more complex but higher data yield sequencing preparation.

## Online Methods

### DNA synthesis

DNA synthesis was performed using standard phosphoramidite chemistry [14] without capping. Volumes and times, described in Table 1, used reagents purchased from Glen Research Corporation. For solid support (PN: ML1-3500-5), we used a BioAutomation 50 nmole scale synthesis column containing controlled porosity glass.

**Table 1:** DNA synthesis reagent parameters. *Only performed as final coupling step to add 5’ phosphate.

| Step | Volume (μL) | Time (s) |
| --- | --- | --- |
| deblock | 600 | 50 |
| Act + {A,C,T,G} (1:1) | 350 | 120 |
| Act + Phos. reagent (1:1)* | 350 | 900 |
| Oxidizer | 750 | 10 |

DNA cleavage was performed in 32% ammonia at room temperature for 1 hour before eluting. Deprotection continued for an additional 11 hours in the same ammonia solution in the storage vessel.

### Sequencing preparation

The extended adapter was constructed from a 1 kilobase fragment that was PCR amplified from the lambda genome using hot start TAQ DNA polymerase (NEB M0496) with a Bsa-I restriction site added by the forward primer. The resulting fragment after digestion had a 3’ A overhang and a 5’-GCGT sticky end on the bottom strand. The fragment was then T/A ligated and prepped according to Oxford Nanopore Technology’s (ONT) LSK-108 kit protocol, yielding the extended adapter with a four base sticky end.

The extended adapter was then mixed according to Table 2 into a sequencing master mix that is used in automated sequencing prep. Thirty minutes prior to sequencing, the master mix was combined with the hairpin oligo and incubated. DTT was left out of the T4 buffer because it damages the nanopores and causes sequencing to fail.

**Table 2:** Sequencing prep master mix. *DTT-free 1× T4 buffer: 50 mM Tris-HCl, 10 mM MgCl_2_, 1mM ATP.

| Reagent | Volume (μL) |
| --- | --- |
| Extended adapter | 15 |
| T4 DNA ligase (NEB: M0202) | 5 |
| DTT-free 10× T4 buffer* | 20 |
| ONT RBF | 93 |
| Nuclease-free water | 64 |
| Total | 197 |

### Nanopore sequencing

Nanopore sequencing was done with an Oxford Nanopore Technologies MinION using an MIN-107 R9.5 flowcell and MinKNOW 18.7.2.0 software. Base calling was performed in 4000 event batches using albacore 2.3.1. The read length distribution and write-to-read quality test were loaded manually (as described in the instructions for LSK-108 sequencing kits); the end-to-end code, write, read, and decode experiment was loaded automatically from the storage vessel.

### Coding and decoding

Data was coded using a two-layer scheme that stored 5 bytes over 32 dsDNA bases with an additional 13 bases of 3’ padding to compensate for lost fidelity near the read end (Fig. 2). The outer layer consisted of a (31,26) Hamming code over a four-symbol alphabet [5] with a checksum base that detects all two-base read errors and corrects all single-base errors. To increase error detection, 6 of the 26 data bases stored a 12-bit hash of the payload, which was checked after decoding to ensure data integrity.

For decoding, groups of 4000 reads were collected and base called using ONT’s Albacore software on 12 CPU cores. Reads that passed QC in Albacore were then aligned to the extended adapter and sequenced for further filtering. Only reads that appeared to have a correctly sized payload region between the adapter sequence and the poly-T hairpin were sent for error checking and decoding.

### DNA alignment

All DNA alignment was done using the parasail `parasail_aligner` command line tool [15] with arguments `-d -t 1 -O SSW -a sg trace striped 16 -o 8 -m NUC.4.4 -e 4`. Alignments to the adapter sequence for decoding used the additional flag `-c 20`, while payload error analysis used flag `-c 8`.

